# Global and national trends in documenting and monitoring species distributions

**DOI:** 10.1101/2020.11.03.367011

**Authors:** Ruth Y. Oliver, Carsten Meyer, Ajay Ranipeta, Kevin Winner, Walter Jetz

## Abstract

Conserving and managing biodiversity in the face of ongoing global change requires sufficient evidence to assess status and trends of species distributions. Here we analyze national trajectories in closing spatiotemporal knowledge gaps for terrestrial vertebrates (1950-2019) based on novel indicators of data coverage and sampling effectiveness. Despite a rapid rise in data coverage, particularly in the last two decades, strong geographic and taxonomic biases persist. For some taxa and regions, a tremendous growth in records failed to directly translate into newfound knowledge due to a sharp decline in sampling effectiveness. But nation’s coverage is stronger for species they hold greater stewardship for. As countries under the post-2020 Global Biodiversity Framework renew their commitments to an improved, rigorous biodiversity knowledge base, our findings highlight opportunities for international collaboration to close critical information gaps.

## Main Text

Detection, understanding, and management of global biodiversity change and its manifold consequences in a rapidly transforming world rely on comprehensive evidence to establish baselines and assess changes. As discussions of the post-2020 Global Biodiversity Framework of the Convention on Biological Diversity (CBD) enter their final stage, the availability of data and metrics to assess progress towards agreed-upon targets has taken a central role (*1*–*5*). The fundamental need for an improved and shared knowledge base of global biodiversity is recognized in the proposed Target 19, which requires the availability of reliable information on biodiversity status and trends.

Descriptions of species’ geographical ranges and their temporal dynamics are fundamental biodiversity measures, as captured in the species distribution Essential Biodiversity Variable (*6*). Status and trends of species’ geographic distributions directly relate to their scope of ecological relevance, population size, and extinction risk, and are thus central to the conservation and management of species and their ecological functions (*7*–*9*). Ambitions to limit threats to species and ensure the integrity of ecosystems, which are central goals of the post-2020 Global Biodiversity Framework under discussion (*10*), critically rely on effective documentation and monitoring of species distributions (*4*, *5*).

Thanks to significant advances in data collection, mobilization, and aggregation (*11*–*13*), publicly accessible occurrence data are growing rapidly, with over 1.6 billion occurrence records across sources and taxa available in the Global Biodiversity Information Facility (GBIF). These data represent an increasing array of sources, including museum specimens, field observations, acoustic and visual sensors, and citizen science efforts (*14*). Digital platforms such as Map of Life (MOL) have begun to integrate these data through models to bolster a multitude of research and conservation applications (*6*, *15*).

Increases in data quantity alone, however, provide little information about overall progress toward an effective spatial biodiversity knowledge base, as records may be highly redundant and cover a limited set of species and regions (*16*). Indeed, prior work has revealed significant taxonomic and geographic gaps in the existing data (*16*–*19*) and highlighted the importance of accounting for expectation and scale in the assessment of coverage (*14*, *16*, *20*–*22*). Scientists have identified a range of socio-economic and ecological drivers for gaps and biases in the current data and identified geographical access, availability of local funding resources, and participation in data-sharing networks as key correlates (*16*).

The aforementioned gaps in knowledge highlight the importance of a more informed and coordinated approach to developing an effective spatial biodiversity evidence base. Developing such an evidence base requires metrics that relate changes in biodiversity data coverage over time to decision-making. As stewards of their biodiversity, nations hold the key to incentivizing an improved information base and stand to gain the greatest benefits from broadly improved biodiversity information by enabling monitoring and robust management decisions. Despite the urgent need to meet international targets and numerous documentations of growing data (*23*, *24*), published work has yet to provide quantitative metrics to track nations’ progress in closing spatiotemporal biodiversity data gaps.

Here, we provide an updatable framework and two national indicators in support of the global assessment, monitoring, and decision-support around annual trends in spatiotemporal biodiversity information. Specifically, we advance and globally implement the Map of Life Species Status Information Index (SSII) which was developed under the auspices of the GEO Biodiversity Observation Network (*25*)(https://mol.org/indicators/coverage) in support of IPBES reporting (https://ipbes.net/core-indicators) and global assessment processes (*10*), as well as the Species Sampling Effectiveness Index (SSEI). We use the indicator framework to compare global and national trends in spatiotemporal biodiversity knowledge since 1950 for over 31,000 terrestrial vertebrate species and over 600 million verified and taxonomically harmonized occurrence records at the level of species, nations, and the globe. We provide a first global assessment for terrestrial vertebrates in this study and infrastructure to continuously track these indices into the future at Map of Life (https://mol.org/indicators/coverage).

The SSII quantifies spatiotemporal biodiversity data coverage for a particular grid resolution and species geographic range expectation (Fig. 1). Global SSII tracks the proportion of expected range cells with records, either for a single species or averaged across multiple species. National SSII restricts this calculation to the range cells inside a particular country. Steward’s SSII follows the National SSII calculation but additionally applies a species-level weight to account for different national stewardships of species. Nations’ varying responsibilities are determined by the portion of a species’ global range they hold (e.g., 1 for country endemics; See Fig. 1 for illustration and Methods for formal description). For a given species, SSII is determined not just by the number of records, but also by how effectively the records cover a species’ full geographic range. We characterize sampling effectiveness by relating the realized spatial distribution of records to the ideal uniform distribution based on Shannon’s entropy (Fig. 1, see Methods) normalized to vary between 0 and 1, a metric we call the Species Sampling Effectiveness Index (SSEI). SSEI has the same properties as SSII and can be calculated at the species, national, or global level and additionally can be adjusted by national stewardship for species.

**Figure 1.**
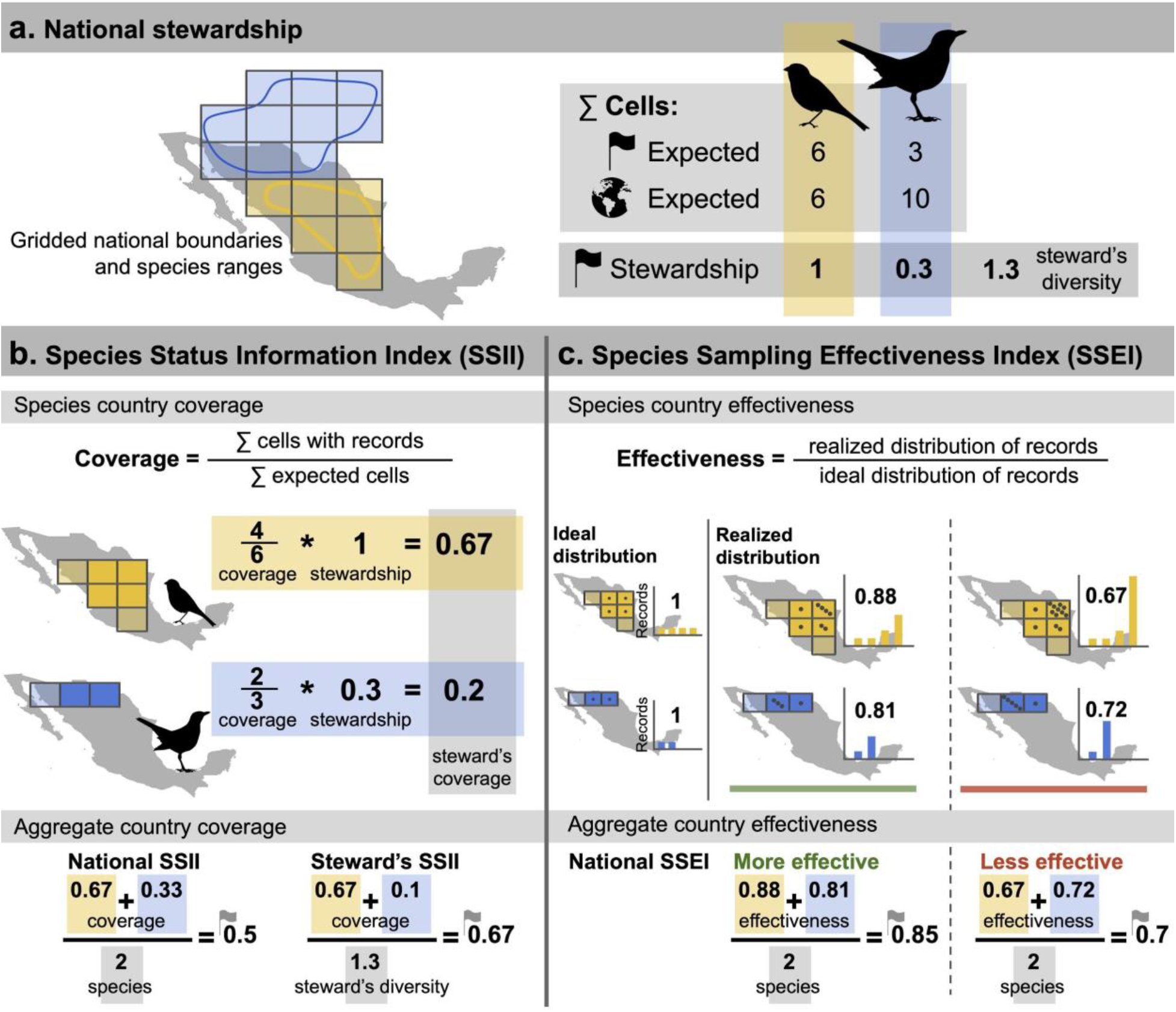
Species Status Information Index (SSII) and Species Sampling Effectiveness Index (SSEI) metrics of biodiversity data coverage and effectiveness. The metrics are illustrated for two hypothetical species with geographic range delineated by binary (e.g., expert range) maps and are assessed for an example 110-km equal-area grid. **a** National stewardship of species is calculated based on the relative portion of species’ ranges falling inside a country. **b** At the species level, the SSII is given as the proportion of cells expected occupied with records in a given year. Species level SSII can be aggregated to the national level via two formulations. National SSII for a given taxonomic group takes the mean across all species expected in a country, Steward’s SSII adjusts the mean across species by their respective national stewardship. **c** Species Sampling Effectiveness Index (SSEI) compares the entropy of the realized distribution of records to that of the ideal distribution (see Methods), where uneven sampling (lower SSEI) is considered less effective than more even sampling (higher SSEI). National SSEI takes the mean across all species expected in a country.

We illustrate the SSII and SSEI for the years 2000-2019 for the jaguar (*Panthera onca*) and collared peccary (*Pecari tajacu*), two widely distributed species with heterogeneous sampling (Fig. 2, Supplementary Table 1). Global SSII was consistently higher for the peccary than the jaguar in the first decade, but the difference was much smaller than the substantial 2-to 10-fold higher annual number of peccary records. Such results suggest a much lower sampling effectiveness, as indexed by the SSEI, for the peccary compared to the jaguar, indicating that many peccary records were concentrated in the same regions. SSII improved markedly for the peccary in recent years, reaching 0.03 (i.e. 3% global range cells with annual records). This increase was associated with increasing SSEI, as the number of records collected were only slightly elevated. National and Steward’s SSII calculated for these two species was highest in Costa Rica and lowest in Brazil (Fig. 2f, Supplementary Table 2). National SSEI was generally highest in Brazil and lowest in Colombia (Fig. 2g).

**Figure 2.**
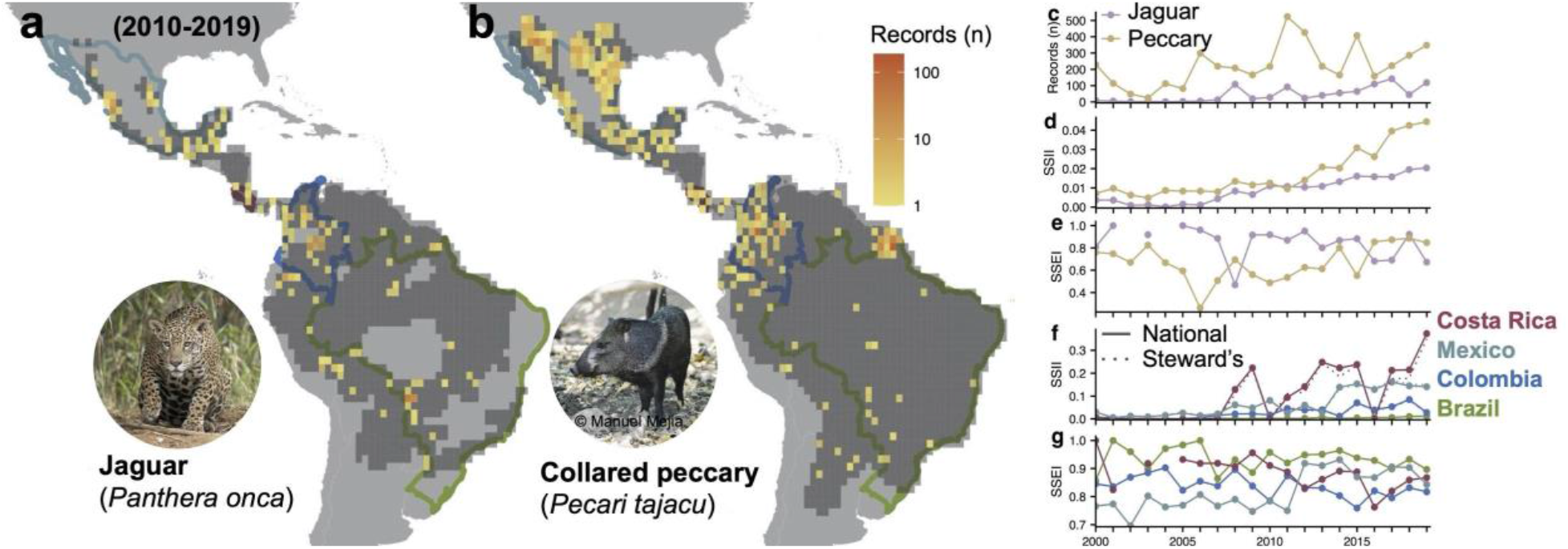
Species and national example patterns and trends. SSII and SSEI trends illustrated for two species, the jaguar (*Panthera onca*) and collared peccary (*Pecari tajacu*). **a-b** The expected occupied cells are shown in dark grey, and total number of records collected 2010-2019 in color. **c-e** Species-level time series of the total number of records (**c**), Global SSII for the whole species range (i.e. all countries with expected range) (**d**), and Global SSEI (**e**) across their expected range. **f-g** Resulting National and Steward’s SSII (**f**) and SSEI (**g**) for four countries.

Biodiversity data collection has rapidly exploded, particularly over the last two decades (Fig. 3a). However, the proliferation of species records and their translation into biodiversity knowledge has played out along substantially different trajectories among taxa. For example, bird species consistently had the largest number of records, with approximately 1,000-fold greater number of records collected annually and 3-fold greater percentage of expected species recorded compared to other terrestrial vertebrates (Fig. 3b). Yet, SSII for birds only exceeded the three other groups after 1980, but has since shown near-linear growth in taxon-wide SSII (Fig. 3c). Coincident with this rapid rise in data collection and coverage for birds species, however, was a rapid decline in sampling effectiveness (Fig. 3d-f). Globally, only approximately half of nations (42%) showed increasing, significant trends in coverage over the previous decade. For those nations showing increases, trends are driven primarily by the rapid increase in avian distribution data (Fig. 4a-c, Supplementary Figure 1, Supplementary Table 3). Interestingly, several world regions that historically most comprehensively sampled the full suite of local species across their geographic ranges are no longer continuing along increasing trajectories. For example, Western Europe, South Africa, and Australia appear to have slowed in their coverage progress, possibly reflecting challenges in the continued mobilization of existing datasets or a lack of impetus to engage in new initiatives (*26*). However, we anticipate that even under constant effort, nations’ coverage may asymptote as marginal gains become more challenging to achieve. Thus, nations with slowing trends may best contribute to CBD goals by partially shifting their investments in national biodiversity data creation toward supporting targeted data mobilization and capacity-building in nations which have so far lagged behind through direct partnerships (*27*). By contrast, much of Asia, South America, and Western and Northern Africa had increasing coverage over the previous decade from initially low values, suggesting encouraging information prospects if trends continue. Our results additionally highlight the importance of regionally targeted capacity-building and data mobilization initiatives such as GBIF’s Biodiversity Information Fund for Asia and Biodiversity Information for Development program focused in sub-Saharan Africa, the Caribbean, and the Pacific.

**Figure 3.**
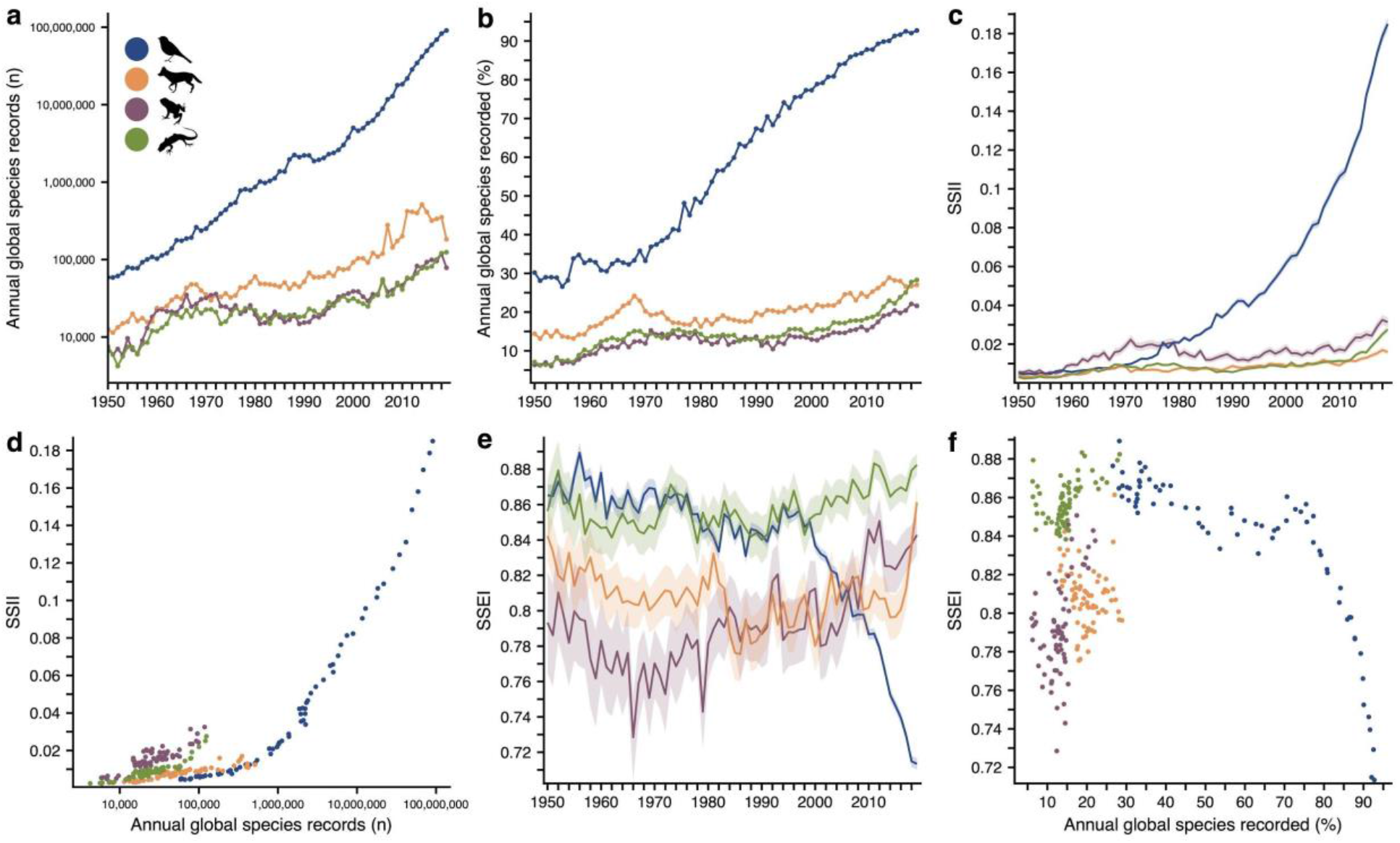
Global trends data coverage and sampling effectiveness across four terrestrial vertebrate groups. **a-c**, Trends in total annual record counts (**a**), percentage of expected species recorded (**b**), and the Global Species Status Information Index (SSII) (**c**). Global SSII is based on data coverage across species’ ranges without consideration of national boundaries. Alternatively, Global SSII for a species is the sum of Steward’s SSII across the nations where it is expected to occur. **d**, Relationship between annual total record counts and Global SSII. **e**, Trends in Global SSEI. **f**, Relationship between percentage of expected species recorded and Global SSEI. **c**,**e** Lines and shading represent means and 95% confidence intervals across species within classes. **d**,**f** Relationships are shown over the past 70 years (1950-2019). Colors in **a-f** indicate birds (blue), mammals (orange), amphibians (purple), reptiles (green).

**Figure 4.**
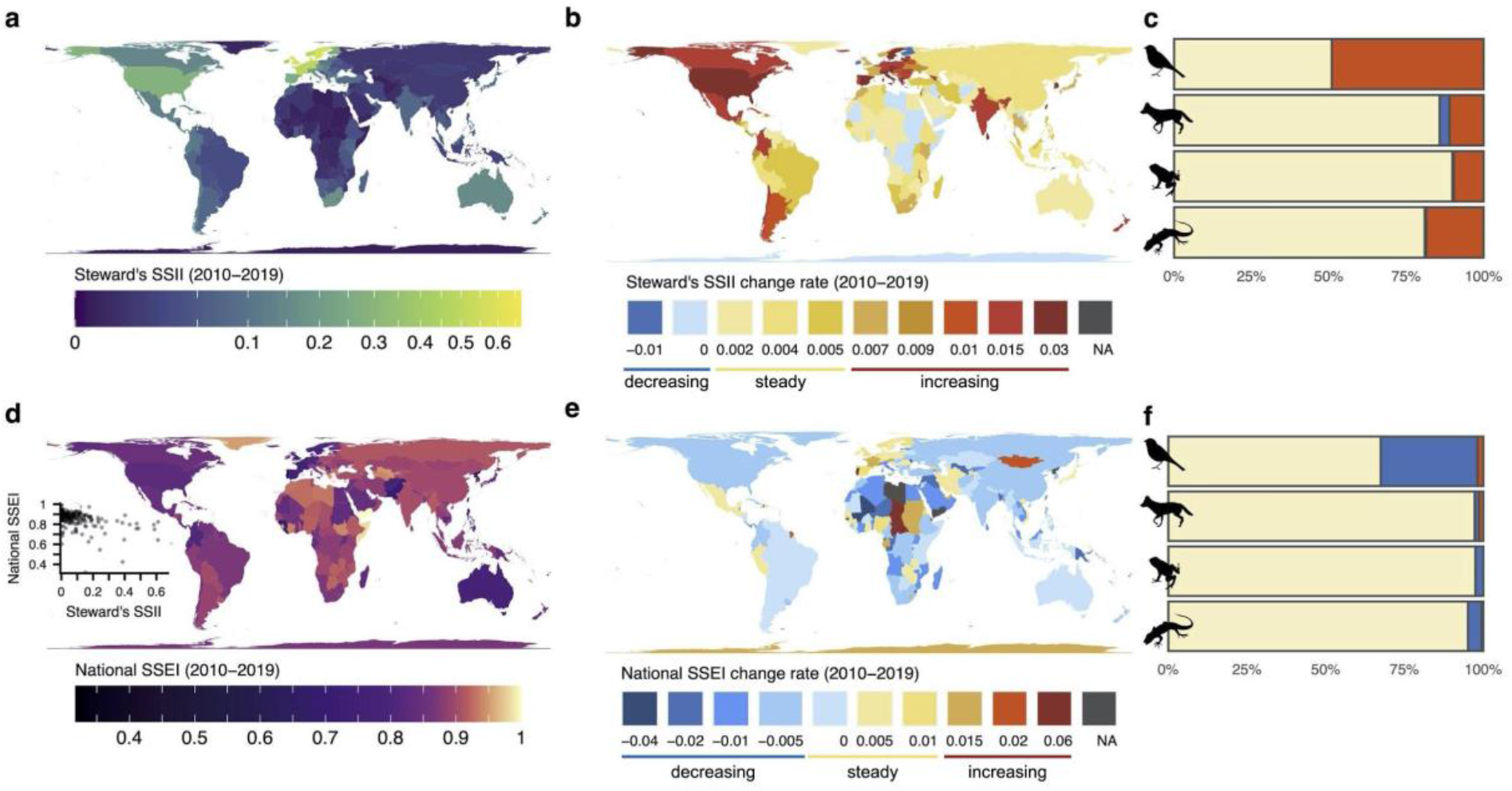
National patterns and trends in spatial biodiversity data coverage and sampling effectiveness. **a,d** Mean Steward’s SSII **(a)** and National SSEI **(d)** over the previous decade (2010-2019) averaged across terrestrial vertebrates; the relationship between data coverage and sampling effectiveness is shown as inset. **b,e** Change rate in Steward’s SSII **(b)** and National SSEI **(e)** over the previous decade. Maximum values for each color bin are labeled below each map. **c,f** Percentage of nations with no significant trends (beige) and significant decreasing (blue) or increasing (red) trends in Steward’s SSII **(c)** and SSEI **(f)** over the previous decade for birds, mammals, amphibians, and reptiles.

While immense in magnitude and potentially useful for other ecological applications, not all data has translated to new knowledge of species distributions (*28*). Sampling effectiveness of biodiversity data as indexed by the SSEI varied considerably among nations over the previous decade, with lowest effectiveness typically within Western Europe, North America, and Australia (Fig. 4d). National trends in effectiveness also appear to be largely driven by the trends in sampling effectiveness for bird species, which has declined rapidly over the past two decades, constraining the direct conversion of the multitudinous accumulation of records into data coverage (Fig. 4e-f).

Differences and tradeoffs in data coverage and effectiveness among taxa appear to be largely driven by the way in which data are collected for different taxonomic groups. Nearly all records for birds in GBIF (>90%) come from direct observations, as opposed to museum specimens which constitute the primary source of records for amphibians, reptiles, and to a lesser degree, mammals (*19*). SSII for birds did not surpass that for other classes until the 1980s, despite having an order of magnitude greater number of records. Further, for the same number of records, birds had the lowest SSII among terrestrial vertebrates and appear to only have achieved the highest SSII through sheer volume of records, as opposed to strategic sampling. Although data coverage for birds increased throughout much of the 20th century, the launch of citizen-science platforms such as eBird (*29*) in the early 2000s undoubtedly played a large role in the expeditious increase in coverage (*14*). However, this onslaught of observations has not been maximally leveraged to enhance the global biodiversity information base, as seen in the coincident decline in avian sampling effectiveness.

The accelerated pace of data coverage for birds compared to other vertebrate groups points to the tremendous role that non-museum based data collection can play in closing knowledge gaps (*30*–*32*). Expanding the impact of citizen science initiatives for information growth will likely benefit from initiatives and guidance addressing the most effective and complementary contributions (*33*, *34*). The rapidly changing landscape of citizen science initiatives will need to be complemented by further supporting and growing coordinated programs through international organisations or government agencies that ensure improved data coverage. Quantifying and identifying particularly important data contributions through products such as the SSII and SSEI, which can be updated and delivered through the Map of Life infrastructure, can support naturalists and initiatives to fill key geographic and taxonomic gaps.

National biodiversity monitoring is influenced by a myriad of social, political, economic, and geographic factors (*16*, *17*, *35*, *36*). We categorized nations into the following four main types based on Steward’s SSII status and trends over the previous decade: (1) coverage below the global mean and no or decreasing trend (42% of nations); (2) increasing coverage, but below the global mean (25%); (3) coverage above the global mean, but no or decreasing trend (16%); (4) both coverage above the global mean and an increasing trend (17%) (Fig. 5a). We highlight national trajectory examples from each group (Fig. 5b). Status and trends in Steward’s SSII differed strongly among continents (Fig. 5c).

**Figure 5.**
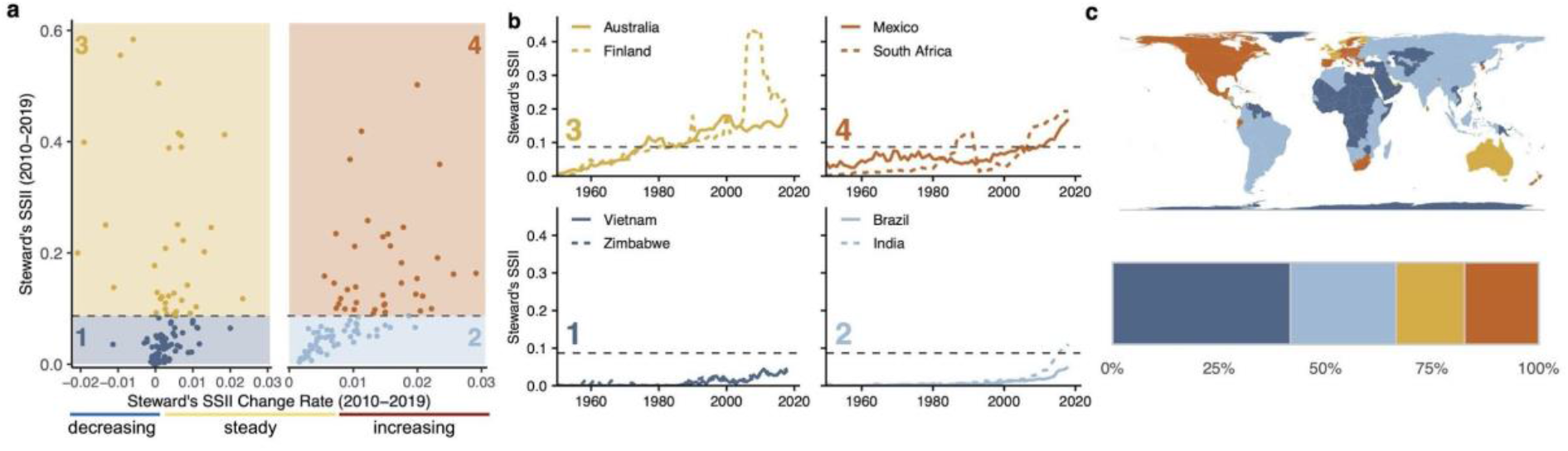
Typologies of nations’ data coverage and trends. **a** Mean values and change rates in Steward’s SSII over the previous decade (2010-2019). Horizontal dashed line represents the global mean of Steward’s SSII. Left panels show nations with no significant or decreasing trends in coverage. Right panels show nations with significant increasing trends in coverage. The four colored quadrants represent nations with: (1) coverage below the global mean and no or decreasing trend; (2) coverage below the global mean and an increasing trend; (3) coverage below the global mean and no or decreasing trend; (4) coverage below the global mean and an increasing trend. **b** Example time series for nations within each type. **c** National assignment to quadrants. Barplot shows percentages of nations within each quadrant.

Political decisions and national infrastructure may have a large influence on data coverage trajectories. For example, the relaxation of stringent data sharing restrictions on biodiversity research in Brazil in 2015 (*37*) appears to have aided in a significant uptick in data coverage. Governmental support for and establishment of a national biodiversity program (CONABIO) (*38*) has placed Mexico on a strong, and increasing, trajectory in both the 20th and 21st centuries. South Africa has achieved particularly high coverage of its biodiversity due to both long-term national policies and large-scale atlasing efforts, such as the Southern African Bird Atlas Project (*39*, *40*).

Through their national commitment to the CBD targets to decrease species extinctions, nations are asked to monitor the species for which they hold greatest responsibility, or, in the case of endemic species, full responsibility. By comparing National and Steward’s SSII, we found that a majority of nations (56%) preferentially survey species for which they hold a high proportion of the global ranges (Supplementary Fig. 2). This may reflect a tendency for endemic biodiversity to confer special cultural importance and for societal interests to influence research agendas (*36*), or simply reflect the preferences of citizen scientists aiming to boost their life lists. Selective monitoring based on nations’ stewardship of species may beneficially promote conservation agendas within nations that have primary control of habitats that species rely on. With this goal in mind, our analysis highlights when nations fall behind on sampling species for which they have high stewardship, and thus play a particularly large role in species’ conservation.

We recognize the limitations of SSII, or any single metric of biodiversity data coverage, to address the range of research and monitoring needs. Our formulation assumes a specific set of taxonomic, spatial, and temporal units and places a burden on nations with particularly high diversity or large national areas to achieve high scores. Furthermore, the annual time units and a relatively coarse grid for the SSII patterns presented here are insensitive to spatiotemporally dense data that could reveal seasonal dynamics and additional insights offered by repeat samples (e.g., in occupancy modelling frameworks) (*41*). In its current form, the SSII also does not account for coverage in environmental space (e.g., as relevant for model-based inference and Essential Biodiversity Variable production; (*6*). Further, the SSII is currently not well suited for rapidly expanding portions of species’ ranges, such as in new invasions, which could be addressed through timely updates of range expectations or other invasion-specific information (*42*).

A group of alternative approaches to quantify sampling completeness rely on parametric or non-parametric richness estimates based on extrapolation of assemblage species accumulation curves (*20*, *43*–*45*). These approaches provide an important complementary contribution especially for extremely under-sampled or - described taxa where globally comprehensive species range expectations, which are necessary for SSII calculation, remain unavailable.

However, results can vary dramatically with specific methodology and structure of input data, resulting in competing recommendations for their development and use (*46*–*48*). The SSII avoids potential pitfalls and limited transparencies of extrapolation approaches by relating record collection directly to best-possible species-level expectations, allowing for decision support at the species level. While this study is limited in scope to terrestrial vertebrates, the framework is easily extended to address other taxonomic groups and realms with ongoing, comprehensive distribution mapping efforts, such as plants and certain marine and invertebrate groups. With the aforementioned limitations and the further potential to extend the metric to account for different spatiotemporal grains, taxa, and data types in mind, the SSII offers an effective initial characterization of biodiversity information at the species, national, and global scales.

## Conclusions

The framework and indicators presented here offer a quantitative and comparable characterization of species, national, and global trajectories in closing biodiversity information gaps. The need for more comprehensive quantitative and standardized biodiversity information to support policy and action not only underpins improved Essential Biodiversity Variables (*6*), but is also recognized as critically needed in recent assessments of IPBES and the post-2020 Global Biodiversity Framework. Our findings suggest that trends in data coverage fundamentally differ by taxa and region and highlight the need to complement and reassess biodiversity sampling strategies to most effectively translate data collection into biodiversity knowledge useful for management and decision-making.

## Acknowledgments

We thank the Map of Life team for their support and expertise, particularly Vijay Barve, Yanina Sica, and Michelle Duong. We are also grateful to the GBIF team, specifically Joe Miller, Tim Hirsch, and Tim Robertson for their help with data and manuscript feedback. This study is supported by the EO Wilson Biodiversity Foundation, NSF grant DEB-1441737 and NASA grants 80NSSC17K0282 and 80NSSC18K0435 to W.J. C.M. acknowledges funding by the Volkswagen Foundation through a Freigeist Fellowship (A118199), and additional support by iDiv, funded by the German Research Foundation (DFG–FZT 118, 202548816).

## Author contributions

R.Y.O., C.M. and W.J. conceived of the study. W.J. conceived the SSII metrics. K.W. contributed to the development of the SSEI metric. R.Y.O. conducted the analyses with contributions from A.R. All authors contributed to the interpretation of the results. R.Y.O. and W.J. led the writing of the manuscript with input from all authors.

## Methods

### Species distribution data

Spatial biodiversity data coverage, hereafter referred to as data coverage, can be calculated at any spatial, temporal, or taxonomic resolution of interest. Doing so requires an estimation of expected biodiversity. We estimated terrestrial vertebrate diversity based on composites of single species distribution maps, which have been shown to approximate species’ global extents over long time periods (ca. 20-40 years)(*49*, *50*), with minimum false presence rates at spatial grains of 100-500 km (*50*). We determined expected diversity for terrestrial birds (N = 9687) (*51*), mammals (N = 5513) (*52*), amphibians (N = 6275) (*53*), and reptiles (N = 9574) (*54*) using a global equal-area grid with the finest spatial grain appropriate (110 x 110 km at the equator). We consider expected diversity of terrestrial vertebrates to be largely static over our study period (1950-2019) with any species range shifts having little impact on the baseline broad-scale, cross-taxon diversity patterns while acknowledging that range expectations likely have higher uncertainty in the earliest decades.

As a representation of digitally accessible and publicly available spatiotemporal biodiversity records, we compiled over one billion occurrence records for terrestrial vertebrates (downloaded April 2020), aggregated by the Global Biodiversity Information Facility (GBIF), of which 661.3 million were taxonomically and spatially valid. To link GBIF-facilitated records to species range maps, we performed taxonomic harmonization based on synonym lists built for this specific purpose. We used scientific names associated with records, which are pre-filtered through GBIF’s backbone taxonomy. Records were considered taxonomically valid if the scientific name could be resolved based on custom-built synonym lists which aggregated synonyms to link to accepted names from additional databases. We followed species delimitations for birds from (*55*), for mammals from (*56*), for amphibians from (*57*), and for reptiles from (*58*). We linked accepted scientific names to potential synonyms and typographical variants compiled from additional data sources (*52*, *53*, *59*–*61*). We restricted our analysis to unambiguous synonyms to avoid matching records to multiple species. We considered records spatially valid based on their intersection with gridded species expert range maps. Restricting records which occurred within expert expectations may exclude true presences, however doing so eliminates errant records as well as those originating from captive or invasive animals.

### Species Status Information Index (SSII)

For a given species, the Species Status Information Index (SSII) captures how well existing data covers the species’ expected range. At the species level, the SSII can be computed across the entirety of the species’ expected range, ignoring national boundaries, or separately within each nation where it is expected to occur. The global SSII,*I_i_* = *O_i_*/*E_i_*, for species *i* across its entire range is the proportion of expected cells, *E_i_*, with records over a given timespan, *O_i_*.

At the national level, for a given taxon with S_c_ species expected in country *C*, we define the SSII, *I_C_*, as follows, distinguishing two formulations (Fig. 1):

1. **National SSII:** This index measures how well, on average, species are documented in a given nation over a given timespane, in this case per year. The index value *I* for country *C* in a particular year is given by the arithmetic mean among expected species *S_c_* of the proportion of expected cells in country *C*, *E_c_*, with records from that year, *O_c_* (Fig. 1b):

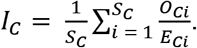
2. **Steward’s SSII:** This index adjusts the national coverage based on nations’ stewardship of species, upweighting the documentation of species for which a nation has particularly high stewardship. National species stewardship *N_Ci_* = *E_Ci_*/*E_.i_* is defined as the proportion of global cells that country *C* holds for a species *i* where *E_.i_* = ∑_*C*_ *E_Ci_* is the total number of expected grid cells for a species across all countries (Fig. 1, top panel). National species stewardship can then be used to weight both the national-level coverage, *I_C_*, and number of species that country *C* is responsible to document, *D*_*steward′s,c*_ (Fig. 1c):

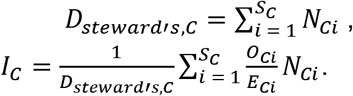

In the current form, the SSII is not capable of capturing true absence information. The expected diversity based on species range maps represents summaries over many years and thus a species is not necessarily expected to occur in every cell in every year due to metapopulation and range dynamics, and differences in suitable environmental conditions. This behavior will certainly depress index values, however, we expect this underestimation to be negligible when averaged across sufficiently large numbers of species constituting a taxon. However, we acknowledge that expectations based on expert range maps may vary among species and regions in their accuracy based on available information.

We calculated the SSII for all extant terrestrial vertebrates both with and without consideration of national boundaries. National boundaries were based on the Database of Global Administrative Areas (GADM version 3.6). National boundaries, species range maps, and spatiotemporal biodiversity records were intersected with a global equal-area grid (110 km) using geohash level 5 representations of each dataset. Geohashes are a public domain geocoding system which encodes geographic coordinates with a unique alphanumeric string in a hierarchical structure and known precision. Geohash level 5 represents geographic space with 5 km bounding boxes at the equator that increase in precision as they approach the poles. Geohash level 5 bounding boxes and the respective proportion contained within each geometry were generated for national boundaries, species ranges, and the equal-area grid. All three datasets were then intersected based on common geohashes. The proportion of each grid cell contained within national boundaries was determined based on summing geohash areas weighted by their proportion within each geometry. Grid cells were weighted within data coverage indices based on their proportion within the nation of interest. Biodiversity records were intersected with national grid cells based on shared geohashes which fell fully inside national boundaries. This restriction may eliminate valid records, but avoids erroneously attributing records to nations due to imprecise spatial intersection.

### Species Sampling Effectiveness Index (SSEI)

Species data coverage is determined not just by count of records but also by their complementarity. To estimate the effectiveness of nations’ biodiversity data collection, we computed the evenness of the spatial distribution of records for each species based on Shannon entropy using the R package DescTools (v0.99.36)(*62*). The Shannon entropy H(X) of a random variable X is an information theoretic metric that measures the expected amount of information or uncertainty in that variable’s distribution (*63*):

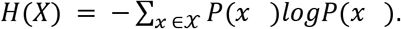

A uniform distribution (*P*(*x*) = 1/|χ| ∀*x* ∈ χ) would have maximum entropy, or evenness, which we denote by *H*\*(*X*) = *log*(|χ|), where |χ| denotes the size of χ, the domain of *X*.

In this application, the entropy *H* of a set of records distributed over *J* cells is given by

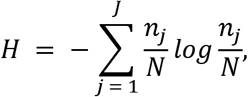

where *n_j_* is the number of records in cell *j* and *N* = ∑_*C*_ *n_j_* is the total number of records. A uniform distribution would represent spatially even sampling of a species, i.e. the same number of records per grid cell and thus *H*\* = *log*(*N*).

We defined the Species Sampling Effectiveness Index (SSEI) for a species as the ratio *H*/*H*\* between the realized evenness of records (i.e. the observed entropy of the distribution of records across all grid cells) and the ideal evenness (i.e. the entropy of a uniform distribution) (Fig. 1c, Supplementary Fig. 3). Similar to SSII, SSEI can be computed at the global, national, or species level and optionally weighted by stewardship at the national level. Global SSEI tracks the ratio of the entropy of the realized and ideal distributions of records for a single species or averaged across many species, without considering national boundaries. National SSEI restricts this calculation to the range cells inside a particular country and takes the average across expected species with data. Steward’s SSEI adjusts the National SSEI based on national stewardship of species, as described for the Steward’s SSII.

### Global trends

We summarized annual trends in the number of occurrence records, the percentage of globally expected species that these records represented, species-level Global SSII and SSEI by class. Statistics were done using R 4.0.0 (*64*) and the package rstatix (v0.6.0) (*65*). We tested for the relationship between the proportion of species recorded and SSEI using Spearman’s rank correlation.

### National trends

We primarily report results for the Steward’s SSII, unless otherwise specified. National data coverage indices were calculated for each terrestrial vertebrate group independently and averaged. Recent Steward’s SSII and SSEI were determined by averaging values of annual SSII and SSEI over the previous ten years (2010-2019). Recent trends in Steward’s SSII and SSEI were determined by testing for significant trends over the previous decade (2010-2019) using linear regression.

We tested for the relationships between variables using Spearman’s rank correlation and differences among classes using ANOVA and Tukey post-hoc test, first testing for normality by Shapiro-Wilk’s test.

We categorized nations into the following four main typologies of status and trends in Steward’s SSII over the previous decade: (1) nations with coverage below the global mean and no or decreasing trend; (2) nations with increasing coverage, but below the global mean; (3) nations with coverage above the global mean, but no or decreasing trend; (4) nations with both coverage above the global mean and an increasing trend.

We estimated nations’ recent propensity to survey local and highly endemic species as compared to those with larger, multinational ranges by calculating the percentage difference between nation’s mean national and Steward’s SSII over the previous decade (2010-2019).

## Supplementary Text

### Global trends

At the global scale, spatiotemporal species records have grown rapidly over the previous 70 years (1950-2019) (Fig. 3a). Bird species consistently had the largest number of records, with approximately 1,000-fold greater number of records collected annually and 3-fold greater percentage of expected species recorded compared to other terrestrial vertebrates (Fig. 3b). The temporal patterns in SSII are different, with birds only exceeding the three other groups after 1980, but since then showing near linear-growth in taxon-wide SSII and exceeding other classes in 2019 by nearly 3-fold (Fig. 3c).

Restricting class-wide SSII values for each class to the years with comparable number of records (58k-118k records; i.e., years 1950s for birds, 1990s to mid 2000s for mammals, and late 2010s for the other two classes), mean SSII was highest for amphibians (0.026), reptiles (0.017), mammals (0.009), and lowest for birds (0.005) (Fig. 3d). SSEI declined by 15% over the previous two decades for bird species and was highest for reptile species for much of the past 70 years (Fig. 3e). Birds had a negative relationship between the percentage of species recorded and sampling effectiveness (Fig. 3f; Spearman’s rho = −0.89, p < 0.001).

### National trends

Over the previous decade, Steward’s SSII varied greatly among nations (Fig. 3a, Supplementary Table 3), with generally higher coverage in Europe, Australia, and the Americas. With the exception of Réunion and Taiwan, all ten nations with the highest data coverage were in Europe.

Steward’s SSII has recently increased in a majority of nations (86%), particularly in North America and southern and eastern Europe with nearly half of nations (42%) showing significant (p < 0.001) increasing trends (Fig. 3b). Of the minority (13%) with decreasing rates, Finland had the most rapid decrease (−0.021 SSII/year). Despite mostly positive trends, much of Africa and Asia saw only negligible increases in indicator values over the last decade, with the exceptions of India, Sri Lanka, and South Korea which showed large increases in data coverage. Nations were nearly evenly split between either non-significant and significantly increasing Steward’s SSII for resident bird species (50.9% and 49.1%, respectively, none decreasing; Fig. 3c). Most nations did not have significant trends in data coverage for mammals (85.8%), amphibians (89.9%), and reptiles (81%).

Recent National SSEI differed strongly among nations (Fig. 4d, Supplementary Table 3). National SSEI was generally lower within western Europe, North America, and Australia. National SSEI and Steward’s SSII were weakly, negatively correlated (Spearman’s rho = −0.43, p < 0.001). A majority of nations (56%) had decreasing SSEI across terrestrial vertebrates, however only 11% of nations globally had significant (p < 0.01), decreasing trends (Fig. 3e). These nations included the United States, Canada, Italy, and South Africa. Decreasing trends in SSEI were most common for bird species (31%) (Fig. 4f).

A majority (63.6%) of nations had increasing Steward’s SSII and National SSEI (Supplementary Fig. 1a). Nations’ mean SSII over the previous decade generally increased with the total number of biodiversity records collected (Supplementary Fig. 1b; Spearman’s rho = 0.55, p < 0.001) and the proportion of expected species recorded (Supplementary Fig. 1c; Spearman’s rho = 0.73, p < 0.001). Mean National SSEI was not significantly related to the proportion of expected species recorded (Supplementary Fig. 1d; Spearman’s rho = −0.23, p = 0.002).

We categorized nations into the following four main types based on Steward’s SSII status and trends over the previous decade: (1) coverage below the global mean and no or decreasing trend (42% of nations); (2) increasing coverage, but below the global mean (25%); (3) coverage above the global mean, but no or decreasing trend (16%); (4) both coverage above the global mean and an increasing trend (17%) (Fig. 4a). We highlight national trajectory examples from each group (Fig. 4b). Status and trends in Steward’s SSII differed strongly among continents (Fig. 4c).

By comparing National and Steward’s SSII, we found that for over half of nations (58.7%), incorporating stewardship increased coverage by over 10%, whereas 26% showed little change (−10 to 10%) (Supplemental Fig. 2). For some countries (14.7%), incorporating stewardship decreased coverage. For example Steward’s SSII was less than half of National SSII for Niger (43.6%) and the Central African Republic (40%), indicating that their endemic or near-endemic species had much lower coverage than other species. In contrast, Steward’s SSII strongly exceeded National SSII in small island nations such as Mayotte (337.9%) and Comoros (184.2%), suggesting their high-stewardship species receive particular recording attention.

---

**Supplementary Figure 1.**
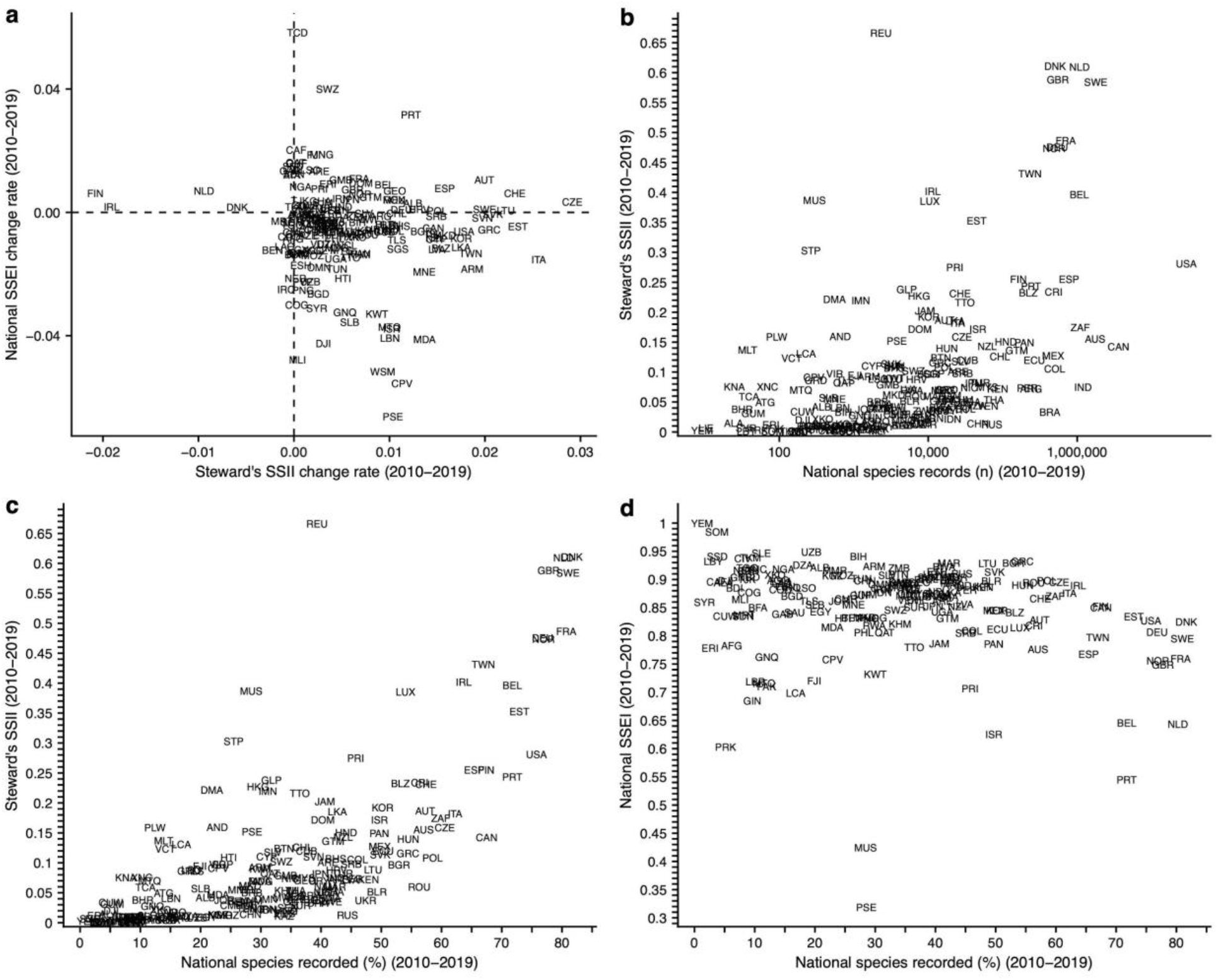
National patterns in data collection, coverage, and sampling effectiveness (2010-2019). **a** Change rates in Steward’s SSII and National SSEI. Dashed lines represent zero slopes. **b-c** Relationship and mismatch between Steward’s SSII and total spatiotemporal records collected nationally (**b**) and the percentage of expected species nationally recorded (**c**). **d** Relationship between the percentage of of expected species nationally recorded and mean National SSEI.

**Supplementary Figure 2.**
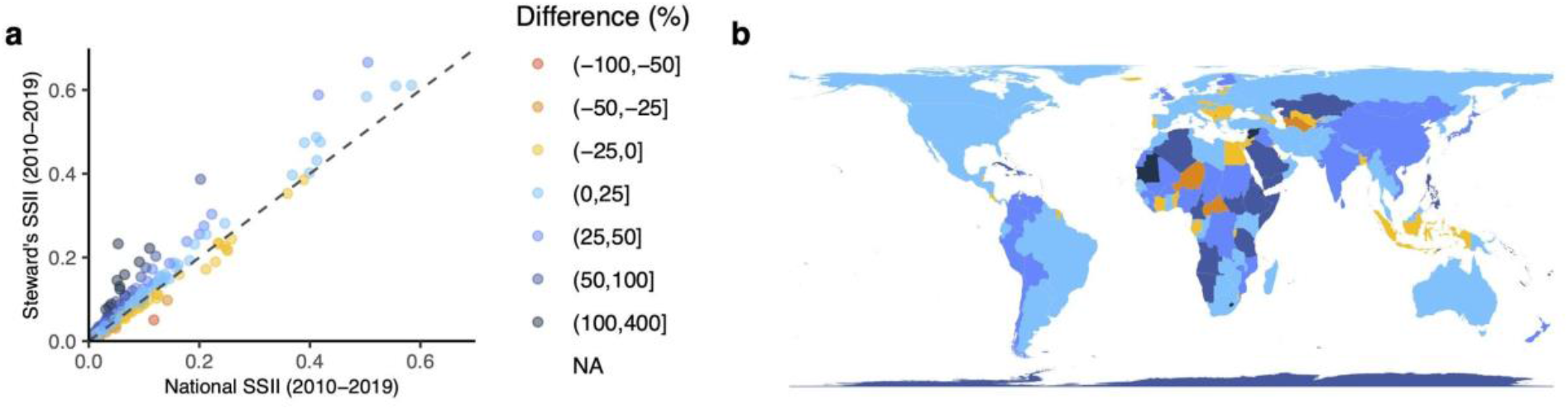
National stewardship in data coverage. **a,** National and Steward’s SSII over the previous decade (2010-2019). Points are colored by the percent difference between national and Steward’s SSII. Dashed line represents the 1:1 line between variables. **b,** Relative stewardship of nations, as estimated by percent difference, over the previous decade. Color scale matches that in panel (**a**).

**Supplementary Figure 3.**
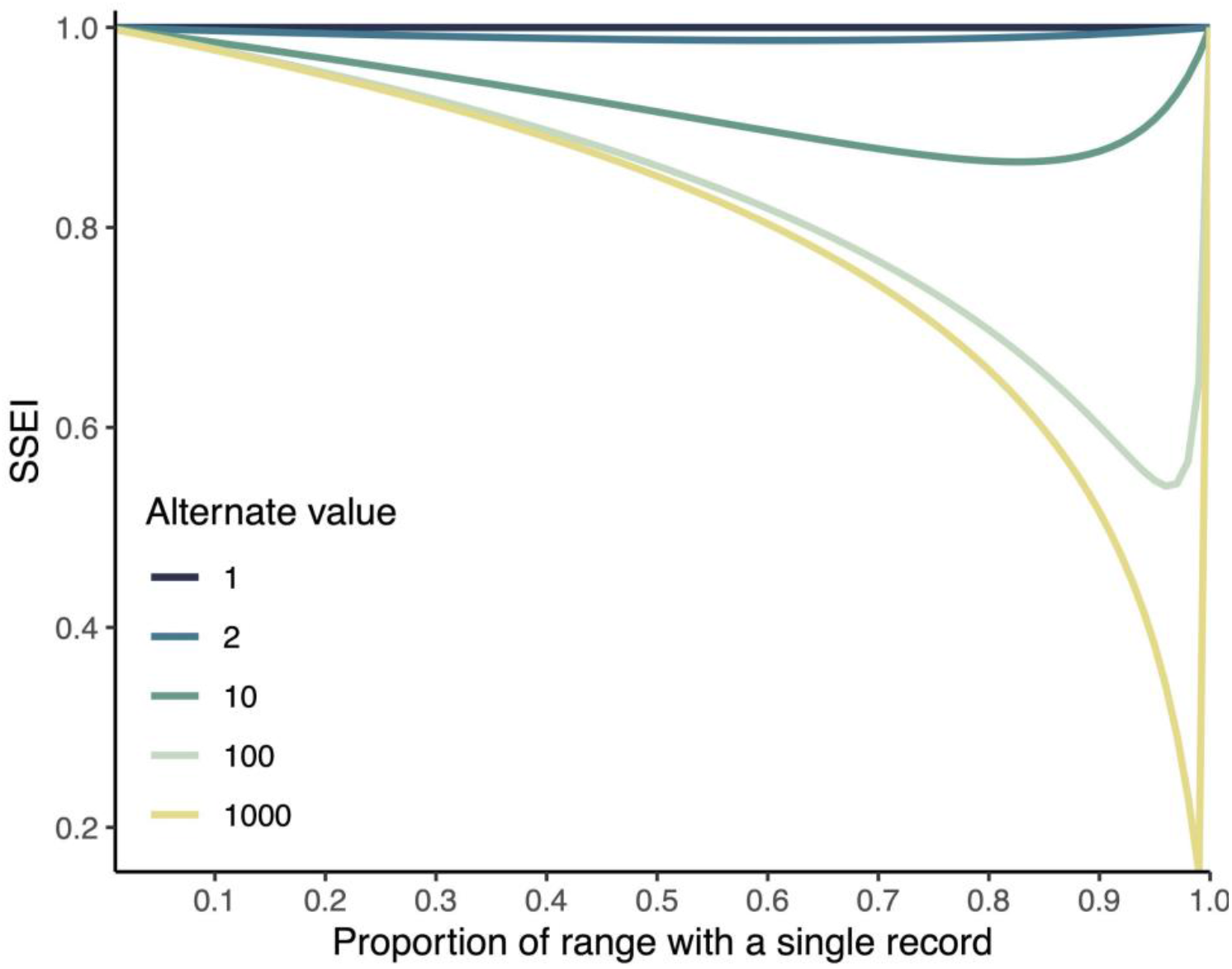
Theoretical examples of the Species Sampling Effectiveness Index (SSEI). Each line corresponds to theoretical cases with different levels of evenness of the distribution of biodiversity records for an idealized species with the same range size. In these examples the proportion of the sampled range with a single record vs. alternate values (1, 2, 10, 100, 1000) is adjusted from 0 to 1. SSEI is highest in cases with uniform or near-uniform sampling (i.e., all grid cells either contain one or two records). SSEI is lowest in cases with highly uneven sampling (i.e., a mixture of grid cells with either a single record or 100-1000 records). These examples also highlight that SSEI is identical in the cases where redundant sampling is uniform (i.e., values are the same if all cells have a 1, 10, or 1000 records). Additionally, SSEI approaches the maximum value when only a small minority of cells contain more than a single record (i.e., the proportion of cells with a single record > 90%).

**Supplementary Table 1.** Species example coverage and sampling effectiveness values. Values presented for the jaguar (*Panthera onca*) and collared peccary (*Pecari tajacu)* as demonstrated in Fig. 2c-e.

| Species | Year | Records (n) | SSII | SSEI |
| --- | --- | --- | --- | --- |
| Panthera onca | 2000 | 9 | 0.004 | 0.810 |
| Panthera onca | 2001 | 5 | 0.004 | 1.000 |
| Panthera onca | 2002 | 1 | 0.001 | NA |
| Panthera onca | 2003 | 3 | 0.001 | 0.918 |
| Panthera onca | 2004 | 1 | 0.000 | NA |
| Panthera onca | 2005 | 3 | 0.002 | 1.000 |
| Panthera onca | 2006 | 5 | 0.001 | 0.961 |
| Panthera onca | 2007 | 13 | 0.004 | 0.885 |
| Panthera onca | 2008 | 108 | 0.008 | 0.469 |
| Panthera onca | 2009 | 20 | 0.007 | 0.916 |
| Panthera onca | 2010 | 28 | 0.011 | 0.918 |
| Panthera onca | 2011 | 92 | 0.011 | 0.869 |
| Panthera onca | 2012 | 23 | 0.010 | 0.951 |
| Panthera onca | 2013 | 40 | 0.011 | 0.802 |
| Panthera onca | 2014 | 55 | 0.013 | 0.868 |
| Panthera onca | 2015 | 65 | 0.016 | 0.883 |
| Panthera onca | 2016 | 110 | 0.016 | 0.681 |
| Panthera onca | 2017 | 142 | 0.016 | 0.691 |
| Panthera onca | 2018 | 45 | 0.020 | 0.922 |
| Panthera onca | 2019 | 119 | 0.020 | 0.673 |
| Pecari tajacu | 2000 | 229 | 0.007 | 0.760 |
| Pecari tajacu | 2001 | 114 | 0.010 | 0.748 |
| Pecari tajacu | 2002 | 48 | 0.006 | 0.670 |
| Pecari tajacu | 2003 | 25 | 0.005 | 0.825 |
| Pecari tajacu | 2004 | 112 | 0.009 | 0.668 |
| Pecari tajacu | 2005 | 82 | 0.008 | 0.594 |
| Pecari tajacu | 2006 | 299 | 0.008 | 0.263 |
| Pecari tajacu | 2007 | 219 | 0.008 | 0.507 |
| Pecari tajacu | 2008 | 208 | 0.013 | 0.694 |
| Pecari tajacu | 2009 | 166 | 0.011 | 0.561 |
| Pecari tajacu | 2010 | 218 | 0.013 | 0.488 |
| Pecari tajacu | 2011 | 524 | 0.009 | 0.535 |
| Pecari tajacu | 2012 | 426 | 0.014 | 0.626 |
| Pecari tajacu | 2013 | 219 | 0.021 | 0.612 |
| Pecari tajacu | 2014 | 165 | 0.020 | 0.801 |
| Pecari tajacu | 2015 | 407 | 0.031 | 0.552 |
| Pecari tajacu | 2016 | 159 | 0.026 | 0.854 |
| Pecari tajacu | 2017 | 222 | 0.040 | 0.873 |
| Pecari tajacu | 2018 | 286 | 0.042 | 0.886 |
| Pecari tajacu | 2019 | 348 | 0.044 | 0.848 |

**Supplementary Table 2.**
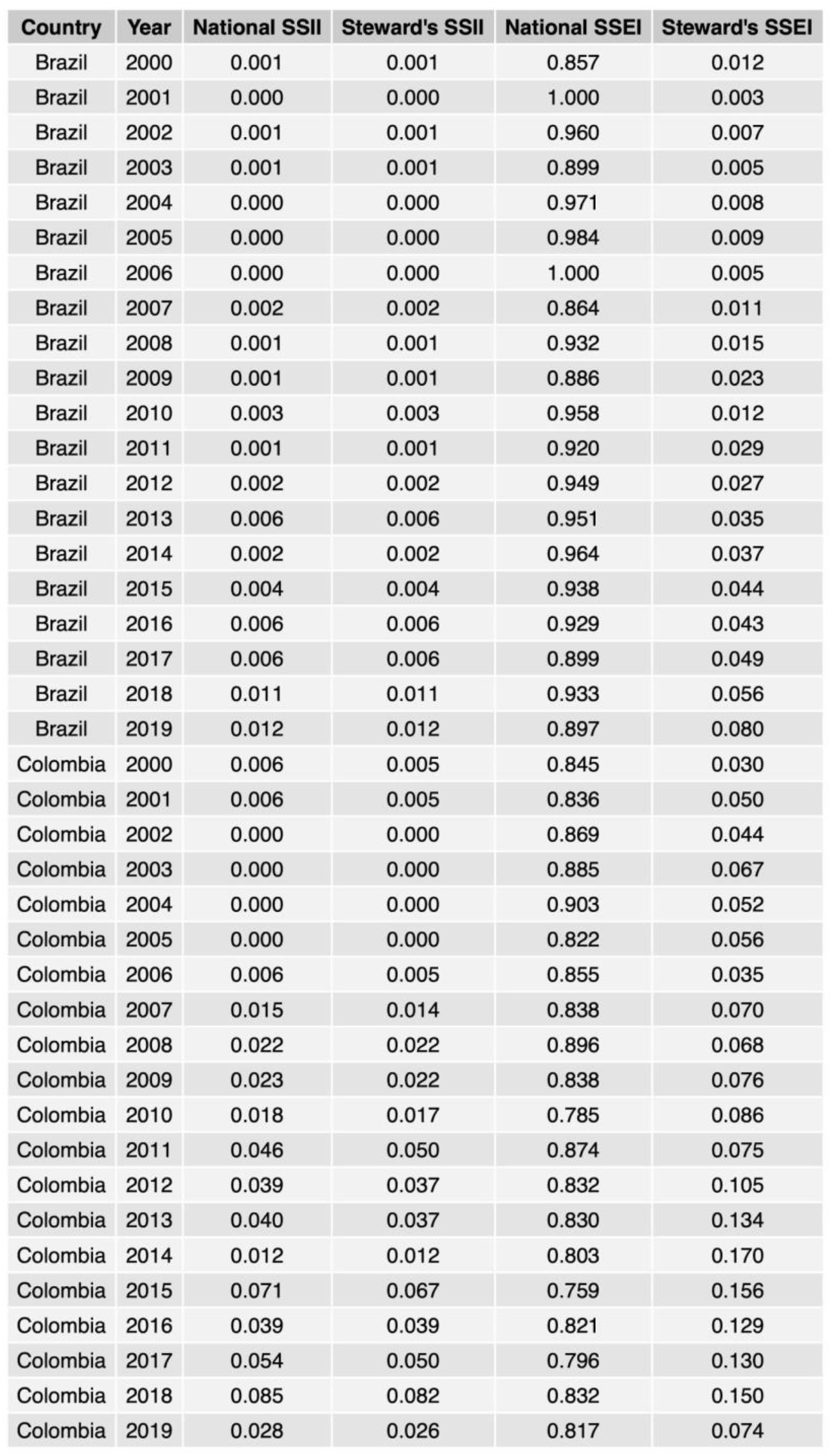
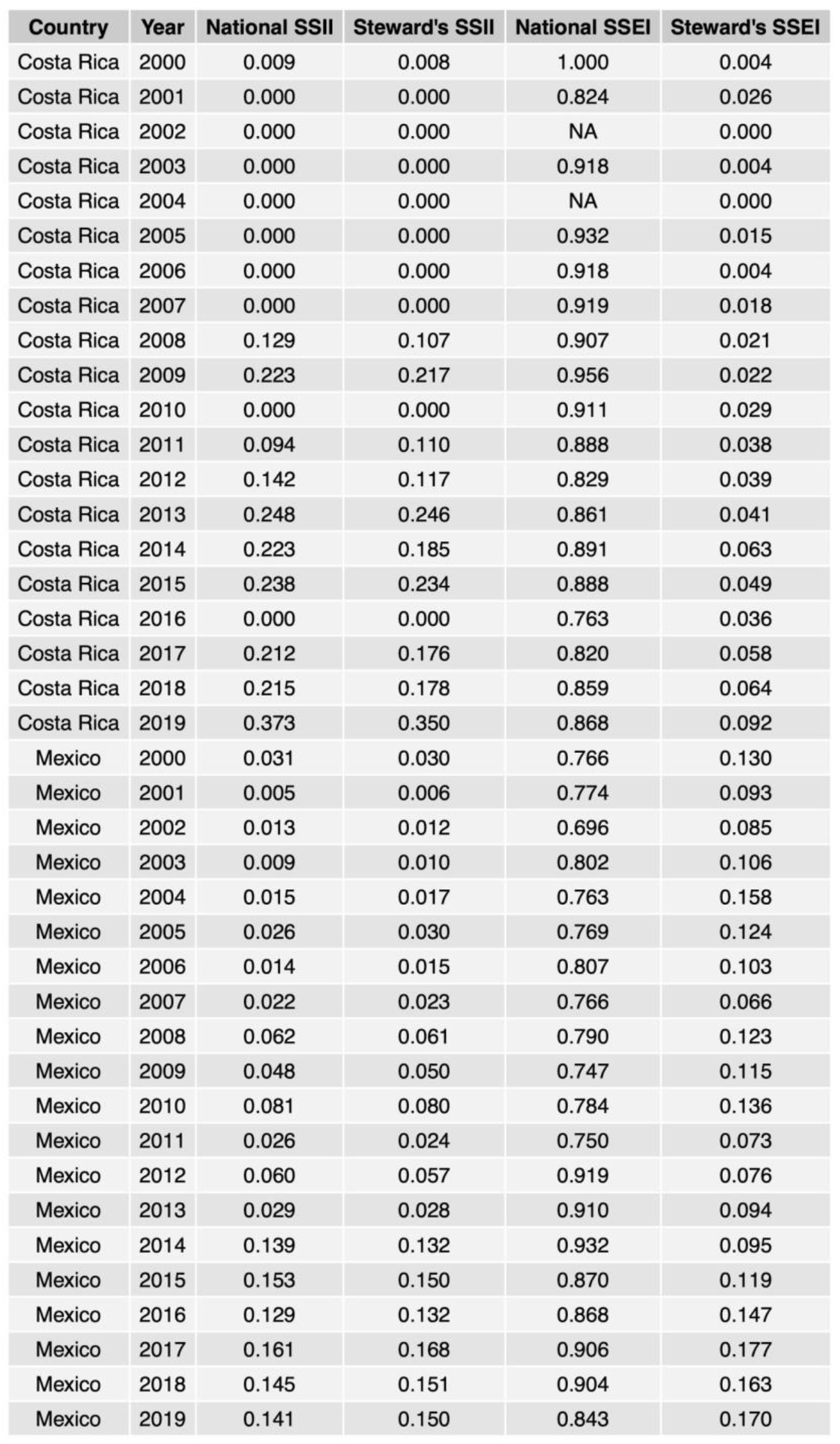
National example data coverage and sampling effectiveness values. Values presented for the jaguar (*Panthera onca*) and collared peccary (*Pecari tajacu)* as demonstrated in Fig. 2f-g.

**Supplementary Table 3.** National data coverage and sampling effectiveness values over the previous decade (2010-2019). ISO3 codes and mean values for National and Stewards’s SSII and SSEI for nations.

| Country | National SSII | Steward's SSII | National SSEI | Steward's SSEI | ISO3 |
| --- | --- | --- | --- | --- | --- |
| Afghanistan | 0.002 | 0.002 | 0.783 | 0.781 | AFG |
| Åland | 0.012 | 0.015 | 0.895 | 0.893 | ALA |
| Albania | 0.051 | 0.042 | 0.921 | 0.915 | ALB |
| Algeria | 0.007 | 0.011 | 0.926 | 0.942 | DZA |
| Andorra | 0.138 | 0.160 | NA | NA | AND |
| Angola | 0.005 | 0.008 | 0.899 | 0.901 | AGO |
| Antarctica | 0.001 | 0.001 | 0.868 | 0.867 | ATA |
| Antigua and Barbuda | 0.051 | 0.050 | NA | NA | ATG |
| Argentina | 0.070 | 0.072 | 0.890 | 0.899 | ARG |
| Armenia | 0.087 | 0.093 | 0.923 | 0.926 | ARM |
| Australia | 0.129 | 0.156 | 0.776 | 0.776 | AUS |
| Austria | 0.154 | 0.187 | 0.828 | 0.818 | AUT |
| Azerbaijan | 0.046 | 0.039 | 0.884 | 0.875 | AZE |
| Bahamas | 0.066 | 0.107 | 0.898 | 0.894 | BHS |
| Bahrain | 0.047 | 0.038 | NA | NA | BHR |
| Bangladesh | 0.014 | 0.011 | 0.871 | 0.866 | BGD |
| Barbados | 0.118 | 0.051 | NA | NA | BRB |
| Belarus | 0.049 | 0.052 | 0.897 | 0.896 | BLR |
| Belgium | 0.368 | 0.397 | 0.646 | 0.655 | BEL |
| Belize | 0.234 | 0.233 | 0.842 | 0.838 | BLZ |
| Benin | 0.040 | 0.036 | 0.832 | 0.849 | BEN |
| Bhutan | 0.098 | 0.124 | 0.909 | 0.911 | BTN |
| Bolivia | 0.028 | 0.039 | 0.901 | 0.895 | BOL |
| Bosnia and Herzegovina | 0.042 | 0.033 | 0.941 | 0.948 | BIH |
| Botswana | 0.034 | 0.041 | 0.923 | 0.925 | BWA |
| Bouvet Island | 0.000 | 0.000 | NA | NA | BVT |
| Brazil | 0.028 | 0.033 | 0.871 | 0.870 | BRA |
| Brunei | 0.019 | 0.009 | 0.915 | 0.923 | BRN |
| Bulgaria | 0.098 | 0.097 | 0.929 | 0.936 | BGR |
| Burkina Faso | 0.003 | 0.005 | 0.850 | 0.841 | BFA |
| Burundi | 0.012 | 0.010 | 0.885 | 0.852 | BDI |
| Cambodia | 0.051 | 0.054 | 0.821 | 0.830 | KHM |
| Cameroon | 0.017 | 0.029 | 0.866 | 0.851 | CMR |
| Canada | 0.124 | 0.143 | 0.851 | 0.855 | CAN |
| Cape Verde | 0.065 | 0.091 | 0.758 | 0.728 | CPV |
| Central African Republic | 0.001 | 0.001 | 0.896 | 0.887 | CAF |
| Chad | 0.002 | 0.002 | 0.905 | 0.911 | TCD |
| Chile | 0.085 | 0.126 | 0.864 | 0.875 | CHL |
| China | 0.010 | 0.014 | 0.898 | 0.899 | CHN |
| Colombia | 0.073 | 0.106 | 0.809 | 0.782 | COL |
| Comoros | 0.051 | 0.145 | 0.888 | 0.828 | COM |
| Costa Rica | 0.235 | 0.234 | 0.818 | 0.801 | CRI |
| Côte d'Ivoire | 0.002 | 0.002 | 0.937 | 0.945 | CIV |
| Croatia | 0.091 | 0.088 | 0.894 | 0.869 | HRV |
| Cuba | 0.079 | 0.120 | 0.870 | 0.856 | CUB |
| Curaçao | 0.017 | 0.034 | 0.835 | 0.842 | CUW |
| Cyprus | 0.093 | 0.110 | 0.884 | 0.879 | CYP |
| Czech Republic | 0.163 | 0.159 | 0.895 | 0.899 | CZE |
| Democratic Republic of the Congo | 0.003 | 0.004 | 0.882 | 0.889 | COD |
| Denmark | 0.584 | 0.611 | 0.825 | 0.831 | DNK |
| Djibouti | 0.009 | 0.020 | 0.898 | 0.892 | DJI |
| Dominica | 0.110 | 0.222 | NA | NA | DMA |
| Dominican Republic | 0.115 | 0.172 | 0.875 | 0.857 | DOM |
| Ecuador | 0.108 | 0.120 | 0.812 | 0.813 | ECU |
| Egypt | 0.010 | 0.010 | 0.843 | 0.845 | EGY |
| El Salvador | 0.101 | 0.118 | 0.907 | 0.903 | SLV |
| Equatorial Guinea | 0.024 | 0.028 | 0.762 | 0.785 | GNQ |
| Eritrea | 0.006 | 0.012 | 0.778 | 0.775 | ERI |
| Estonia | 0.359 | 0.353 | 0.834 | 0.835 | EST |
| Ethiopia | 0.022 | 0.037 | 0.914 | 0.888 | ETH |
| Falkland Islands | 0.022 | 0.033 | 0.886 | 0.857 | FLK |
| Faroe Islands | 0.066 | 0.055 | 0.849 | 0.841 | FRO |
| Fiji | 0.049 | 0.094 | 0.720 | 0.646 | FJI |
| Finland | 0.200 | 0.255 | 0.852 | 0.852 | FIN |
| France | 0.412 | 0.487 | 0.760 | 0.768 | FRA |
| French Guiana | 0.032 | 0.030 | 0.872 | 0.883 | GUF |
| French Polynesia | 0.083 | 0.097 | 0.870 | 0.828 | PYF |
| French Southern Territories | 0.014 | 0.026 | 0.820 | 0.820 | ATF |
| Gabon | 0.011 | 0.010 | 0.839 | 0.846 | GAB |
| Gambia | 0.089 | 0.079 | 0.895 | 0.884 | GMB |
| Georgia | 0.072 | 0.071 | 0.895 | 0.879 | GEO |
| Germany | 0.419 | 0.476 | 0.806 | 0.806 | DEU |
| Ghana | 0.046 | 0.052 | 0.869 | 0.866 | GHA |
| Greece | 0.096 | 0.116 | 0.932 | 0.928 | GRC |
| Greenland | 0.002 | 0.002 | 0.952 | 0.940 | GRL |
| Grenada | 0.074 | 0.086 | NA | NA | GRD |
| Guadeloupe | 0.177 | 0.238 | NA | NA | GLP |
| Guam | 0.048 | 0.031 | NA | NA | GUM |
| Guatemala | 0.118 | 0.136 | 0.832 | 0.815 | GTM |
| Guinea | 0.005 | 0.006 | 0.685 | 0.670 | GIN |
| Guinea-Bissau | 0.009 | 0.010 | 0.904 | 0.937 | GNB |
| Guyana | 0.032 | 0.035 | 0.875 | 0.864 | GUY |
| Haiti | 0.065 | 0.109 | 0.831 | 0.827 | HTI |
| Heard Island and McDonald Islands | 0.036 | 0.037 | NA | NA | HMD |
| Honduras | 0.128 | 0.151 | 0.889 | 0.862 | HND |
| Hong Kong | 0.246 | 0.227 | NA | NA | HKG |
| Hungary | 0.134 | 0.140 | 0.890 | 0.896 | HUN |
| Iceland | 0.229 | 0.189 | 0.848 | 0.858 | ISL |
| India | 0.059 | 0.075 | 0.904 | 0.900 | IND |
| Indonesia | 0.023 | 0.021 | 0.877 | 0.875 | IDN |
| Iran | 0.022 | 0.023 | 0.882 | 0.869 | IRN |
| Iraq | 0.012 | 0.018 | 0.886 | 0.894 | IRQ |
| Ireland | 0.399 | 0.402 | 0.890 | 0.893 | IRL |
| Isle of Man | 0.250 | 0.219 | NA | NA | IMN |
| Israel | 0.212 | 0.172 | 0.625 | 0.634 | ISR |
| Italy | 0.162 | 0.182 | 0.875 | 0.875 | ITA |
| Jamaica | 0.121 | 0.203 | 0.786 | 0.775 | JAM |
| Japan | 0.062 | 0.081 | 0.852 | 0.835 | JPN |
| Jordan | 0.039 | 0.038 | 0.862 | 0.863 | JOR |
| Kazakhstan | 0.007 | 0.011 | 0.895 | 0.897 | KAZ |
| Kenya | 0.061 | 0.072 | 0.887 | 0.877 | KEN |
| Kiribati | 0.000 | 0.000 | NA | NA | KIR |
| Kosovo | 0.031 | 0.022 | 0.908 | 0.912 | XKO |
| Kuwait | 0.099 | 0.090 | 0.732 | 0.725 | KWT |
| Kyrgyzstan | 0.014 | 0.014 | 0.907 | 0.909 | KGZ |
| Laos | 0.012 | 0.018 | 0.894 | 0.895 | LAO |
| Latvia | 0.070 | 0.072 | 0.856 | 0.857 | LVA |
| Lebanon | 0.040 | 0.041 | 0.888 | 0.891 | LBN |
| Lesotho | 0.041 | 0.088 | 0.885 | 0.886 | LSO |
| Liberia | 0.008 | 0.011 | 0.719 | 0.742 | LBR |
| Libya | 0.001 | 0.001 | 0.932 | 0.935 | LBY |
| Liechtenstein | 0.008 | 0.007 | NA | NA | LIE |
| Lithuania | 0.100 | 0.089 | 0.929 | 0.927 | LTU |
| Luxembourg | 0.389 | 0.385 | 0.814 | 0.812 | LUX |
| Macedonia | 0.066 | 0.062 | 0.833 | 0.863 | MKD |
| Madagascar | 0.057 | 0.069 | 0.832 | 0.827 | MDG |
| Malawi | 0.036 | 0.042 | 0.886 | 0.884 | MWI |
| Malaysia | 0.068 | 0.075 | 0.875 | 0.867 | MYS |
| Mali | 0.001 | 0.002 | 0.865 | 0.875 | MLI |
| Malta | 0.118 | 0.137 | NA | NA | MLT |
| Martinique | 0.078 | 0.070 | 0.716 | 0.719 | MTQ |
| Mauritania | 0.004 | 0.010 | 0.837 | 0.819 | MRT |
| Mauritius | 0.202 | 0.387 | 0.425 | 0.361 | MUS |
| Mayotte | 0.053 | 0.233 | NA | NA | MYT |
| Mexico | 0.111 | 0.128 | 0.845 | 0.836 | MEX |
| Micronesia | 0.036 | 0.082 | NA | NA | FSM |
| Moldova | 0.051 | 0.047 | 0.815 | 0.795 | MDA |
| Mongolia | 0.014 | 0.020 | 0.893 | 0.887 | MNG |
| Montenegro | 0.062 | 0.055 | 0.855 | 0.865 | MNE |
| Morocco | 0.050 | 0.060 | 0.930 | 0.922 | MAR |
| Mozambique | 0.009 | 0.013 | 0.908 | 0.908 | MOZ |
| Myanmar | 0.011 | 0.013 | 0.916 | 0.910 | MMR |
| Namibia | 0.034 | 0.061 | 0.871 | 0.880 | NAM |
| Nepal | 0.048 | 0.051 | 0.879 | 0.897 | NPL |
| Netherlands | 0.555 | 0.610 | 0.643 | 0.635 | NLD |
| New Caledonia | 0.094 | 0.153 | 0.828 | 0.824 | NCL |
| New Zealand | 0.106 | 0.143 | 0.852 | 0.843 | NZL |
| Nicaragua | 0.064 | 0.074 | 0.871 | 0.866 | NIC |
| Niger | 0.003 | 0.002 | 0.916 | 0.918 | NER |
| Nigeria | 0.004 | 0.005 | 0.919 | 0.920 | NGA |
| Niue | 0.021 | 0.026 | NA | NA | NIU |
| North Korea | 0.003 | 0.003 | 0.603 | 0.615 | PRK |
| Northern Cyprus | 0.046 | 0.075 | 0.918 | 0.918 | XNC |
| Norway | 0.390 | 0.474 | 0.756 | 0.765 | NOR |
| Oman | 0.036 | 0.040 | 0.892 | 0.883 | OMN |
| Pakistan | 0.004 | 0.005 | 0.710 | 0.705 | PAK |
| Palau | 0.065 | 0.159 | NA | NA | PLW |
| Palestina | 0.139 | 0.152 | 0.318 | 0.314 | PSE |
| Panama | 0.146 | 0.149 | 0.786 | 0.774 | PAN |
| Papua New Guinea | 0.019 | 0.023 | 0.832 | 0.831 | PNG |
| Paraguay | 0.042 | 0.045 | 0.900 | 0.904 | PRY |
| Peru | 0.057 | 0.074 | 0.881 | 0.871 | PER |
| Philippines | 0.036 | 0.055 | 0.806 | 0.795 | PHL |
| Poland | 0.094 | 0.108 | 0.898 | 0.894 | POL |
| Portugal | 0.258 | 0.244 | 0.545 | 0.568 | PRT |
| Puerto Rico | 0.208 | 0.275 | 0.707 | 0.690 | PRI |
| Qatar | 0.093 | 0.083 | 0.806 | 0.844 | QAT |
| Republic of Congo | 0.003 | 0.004 | 0.877 | 0.886 | COG |
| Reunion | 0.505 | 0.666 | NA | NA | REU |
| Romania | 0.062 | 0.060 | 0.895 | 0.892 | ROU |
| Russia | 0.012 | 0.013 | 0.911 | 0.905 | RUS |
| Rwanda | 0.076 | 0.069 | 0.818 | 0.820 | RWA |
| Saint Helena | 0.011 | 0.010 | NA | NA | SHN |
| Saint Kitts and Nevis | 0.031 | 0.076 | NA | NA | KNA |
| Saint Lucia | 0.056 | 0.131 | 0.699 | 0.648 | LCA |
| Saint Pierre and Miquelon | 0.013 | 0.012 | NA | NA | SPM |
| Saint Vincent and the Grenadines | 0.056 | 0.124 | NA | NA | VCT |
| Samoa | 0.091 | 0.189 | 0.669 | 0.682 | WSM |
| São Tomé and Príncipe | 0.222 | 0.303 | NA | NA | STP |
| Saudi Arabia | 0.005 | 0.009 | 0.842 | 0.841 | SAU |
| Senegal | 0.023 | 0.025 | 0.890 | 0.885 | SEN |
| Serbia | 0.108 | 0.098 | 0.805 | 0.806 | SRB |
| Sierra Leone | 0.010 | 0.012 | 0.947 | 0.947 | SLE |
| Singapore | 0.142 | 0.098 | NA | NA | SGP |
| Slovakia | 0.122 | 0.114 | 0.913 | 0.921 | SVK |
| Slovenia | 0.126 | 0.110 | 0.904 | 0.905 | SVN |
| Solomon Islands | 0.034 | 0.057 | 0.855 | 0.858 | SLB |
| Somalia | 0.001 | 0.001 | 0.985 | 0.985 | SOM |
| South Africa | 0.158 | 0.174 | 0.871 | 0.876 | ZAF |
| South Georgia and the South Sandwich Islands | 0.103 | 0.175 | 0.973 | 0.963 | SGS |
| South Korea | 0.182 | 0.192 | 0.845 | 0.860 | KOR |
| South Sudan | 0.001 | 0.001 | 0.941 | 0.950 | SSD |
| Spain | 0.212 | 0.255 | 0.768 | 0.772 | ESP |
| Sri Lanka | 0.146 | 0.186 | 0.877 | 0.840 | LKA |
| Sudan | 0.001 | 0.001 | 0.835 | 0.851 | SDN |
| Suriname | 0.027 | 0.029 | 0.851 | 0.860 | SUR |
| Svalbard and Jan Mayen | 0.090 | 0.092 | 0.744 | 0.693 | SJM |
| Swaziland | 0.123 | 0.102 | 0.846 | 0.854 | SWZ |
| Sweden | 0.502 | 0.584 | 0.795 | 0.821 | SWE |
| Switzerland | 0.191 | 0.232 | 0.866 | 0.848 | CHE |
| Syria | 0.003 | 0.006 | 0.860 | 0.875 | SYR |
| Taiwan | 0.413 | 0.432 | 0.798 | 0.813 | TWN |
| Tajikistan | 0.004 | 0.005 | 0.901 | 0.897 | TJK |
| Tanzania | 0.029 | 0.044 | 0.892 | 0.885 | TZA |
| Thailand | 0.048 | 0.054 | 0.883 | 0.877 | THA |
| Timor-Leste | 0.055 | 0.087 | 0.862 | 0.844 | TLS |
| Togo | 0.009 | 0.008 | 0.920 | 0.920 | TGO |
| Tonga | 0.003 | 0.000 | NA | NA | TON |
| Trinidad and Tobago | 0.251 | 0.216 | 0.780 | 0.795 | TTO |
| Tunisia | 0.024 | 0.025 | 0.902 | 0.879 | TUN |
| Turkey | 0.073 | 0.082 | 0.904 | 0.904 | TUR |
| Turkmenistan | 0.002 | 0.001 | 0.939 | 0.941 | TKM |
| Turks and Caicos Islands | 0.038 | 0.059 | NA | NA | TCA |
| Uganda | 0.047 | 0.050 | 0.843 | 0.829 | UGA |
| Ukraine | 0.033 | 0.037 | 0.888 | 0.893 | UKR |
| United Arab Emirates | 0.099 | 0.101 | 0.862 | 0.863 | ARE |
| United Kingdom | 0.415 | 0.588 | 0.749 | 0.729 | GBR |
| United States | 0.246 | 0.281 | 0.827 | 0.839 | USA |
| Uruguay | 0.060 | 0.070 | 0.909 | 0.909 | URY |
| Uzbekistan | 0.010 | 0.008 | 0.949 | 0.947 | UZB |
| Vanuatu | 0.029 | 0.047 | 0.914 | 0.916 | VUT |
| Venezuela | 0.030 | 0.043 | 0.863 | 0.849 | VEN |
| Vietnam | 0.028 | 0.036 | 0.873 | 0.853 | VNM |
| Virgin Islands, U.S. | 0.087 | 0.097 | NA | NA | VIR |
| Western Sahara | 0.007 | 0.010 | 0.897 | 0.899 | ESH |
| Yemen | 0.001 | 0.002 | 1.000 | 1.000 | YEM |
| Zambia | 0.014 | 0.015 | 0.920 | 0.921 | ZMB |
| Zimbabwe | 0.031 | 0.035 | 0.905 | 0.903 | ZWE |

